# Topological stress is responsible for the detrimental outcomes of head-on replication-transcription conflicts

**DOI:** 10.1101/691188

**Authors:** Kevin S. Lang, Houra Merrikh

**Author notes:** Corresponding author and lead contact Houra Merrikh.

## Abstract

Conflicts between the replication and transcription machineries have profound effects on chromosome duplication, genome organization, as well as evolution across species. Head-on conflicts (lagging strand genes) are significantly more detrimental than co-directional conflicts (leading strand genes). The source of this fundamental difference is unknown. Here, we report that topological stress underlies this difference. We find that head-on conflict resolution requires the relaxation of positive supercoils DNA gyrase and Topo IV. Interestingly, we find that after positive supercoil resolution, gyrase introduces excessive negative supercoils at head-on conflict regions, driving pervasive R-loop formation. The formation of these R-Loops through gyrase activity is most likely caused by the diffusion of negative supercoils through RNA polymerase spinning. Altogether, our results address a longstanding question regarding replication-transcription conflicts by revealing the fundamental mechanistic difference between the two types of encounters.

## Introduction

Transcription and DNA replication occur simultaneously on the same template. The lack of spatiotemporal separation between these two processes leads to conflicts between them every replication cycle. Replication and transcription machineries can encounter each other either head-on or co-directionally. Co-directional conflicts occur when genes are transcribed on the leading strand whereas head-on conflicts occur when genes are transcribed on the lagging strand. It has been demonstrated that head-on conflicts are more deleterious than co-directional conflicts in that they cause increased mutagenesis, DNA breaks, replisome stalling and restart, (Lang et al. 2017; Paul et al. 2013; Million-Weaver et al. 2015; Million-Weaver, Samadpour, and Merrikh 2015; Mirkin and Mirkin 2005; Prado and Aguilera 2005; French 1992; J. D. Wang, Berkmen, and Grossman 2007; C. N. Merrikh and Merrikh 2018; H. Merrikh et al. 2011; Pomerantz and O’Donnell 2010; Hamperl et al. 2017). Despite many insightful studies into these inevitable encounters, the fundamental question regarding why head-on conflicts are more detrimental than co-directional conflicts remains unanswered. It is perplexing that encounters between the same two machineries (the replication machinery or the replisome, and RNA polymerase or RNAP) can have such different outcomes simply due to orientation.

Topological constraints could explain why head-on conflicts are more deleterious than co-directional conflicts. Unwinding of DNA during transcription generates positively supercoiled DNA ahead, and negatively supercoiled DNA behind RNAP (Wu et al. 1988; Liu and Wang 1987). Similarly, during replication, positive supercoils accumulate in front of the replisome (Vos et al. 2011; Postow, Peter, and Cozzarelli 1999; Hiasa and Marians 1996). The resolution of this supercoiled DNA is critical for both transcription and replication to proceed efficiently (Khodursky et al. 2000). In a co-directional conflict, the positive supercoiling generated in front of the replisome would encounter the negative supercoiling produced from active RNAPs ahead. This would most likely cause a net neutral change in local supercoiling levels. However, during a head-on conflict, the positive supercoiling generated ahead of the replisome would encounter the positive supercoiling produced by RNAP. Therefore, in a head-on conflict, there may be a transient buildup of positive supercoils that has the potential to change the fundamental mechanics of the replisome and RNAP. Such changes could stall the replisome, leading to disassembly, and changing the dynamics of RNAP and associated mRNAs. These predictions suggest that torsional stress could be the key driver of conflict severity and therefore this model must be tested.

Another key question is whether topoisomerases are critical conflict resolution factors. The resolution of supercoils in all organisms requires topoisomerases (Champoux 2001; J. C. Wang 2002; Vos et al. 2011). In bacteria, there are two topoisomerases that relax positive supercoils: DNA gyrase and Topo IV. DNA gyrase and Topo IV are both required for replication fork progression *in vivo* (Khodursky et al. 2000; Crisona et al. 2000; Peng and Marians 1993; Ashley *et al*. 2017; Vos et al. 2011). Topo IV also plays a critical role in the resolution of catenanes (intertwined chromosomes) as well as the separation of sister chromatids during segregation (Hiasa and Marians 1996; Zechiedrich and Cozzarelli 1995). If the torsional stress hypothesis is correct, then type II topoisomerases should be critical conflict resolution factors, yet, this question has not been addressed.

Here, we report that type II topoisomerases preferentially associate with head-on genes and that cells harboring engineered head-on conflicts are sensitized to type II topoisomerase inhibitors. Accordingly, we find that conditional depletion of either gyrase or Topo IV is deleterious to cells experiencing engineered head-on conflicts. Inhibition of type II topoisomerase activity leads to increased stalling of the replisome when it approaches a gene transcribed in the head-on, but not the co-directional orientation. Remarkably, however, we find that negative supercoil introduction by DNA gyrase at head-on conflict regions is responsible for the formation of toxic R-loops at these regions. Consistent with this finding, we observe that, in cells lacking the RNase HIII enzyme, which resolves R-Loops, inhibition of type II topoisomerases lowers R-loop abundance, and alleviates R-Loop induced replisome stalling at head-on genes. Furthermore, an allele of gyrase that is strongly defective in introduction of negative supercoils completely rescues the lethality of cells lacking RNase HIII that are experiencing head-on conflicts. This rescue is observed in experiments examining both engineered and endogenous head-on conflicts, which arise predominantly during the expression of stress response genes.

## Results

### Type II topoisomerases preferentially associate with a head-on but not a co-directional engineered conflict region

The relaxation of both positive and negative supercoils is an essential process in all cells. In *B. subtilis*, relaxation of positive supercoils is accomplished by the activity of either gyrase or Topo IV (Vos et al. 2011; Postow, Crisona, et al. 2001; Crisona et al. 2000; Ashley et al. 2017). If the model of positive supercoil accumulation at head-on conflict regions is correct, then these enzymes should preferentially associate with a head-on conflict region. To test this hypothesis, we measured gyrase and Topo IV enrichment genome-wide, using chromatin immunoprecipitation followed by deep sequencing (ChIP-Seq). Most head-on genes are not expressed under standard laboratory conditions. Rather, the majority of head-on genes are induced under specific conditions, such as during exposure to environmental stress (Nicolas et al. 2012; Mostertz et al. 2004; Guariglia-Oropeza and Helmann 2011; Lang et al. 2017). Therefore, we did not expect to see enrichment of type II topoisomerases at endogenous head-on genes during growth in rich media. In order to study the effects of topology at head-on conflict regions, we took advantage of several different tightly controlled engineered conflict systems, all of which were integrated onto the chromosome. In each of these systems the same exact gene (e.g. *lacZ*) was inserted onto the chromosome in the same locus, in either the head-on or co-directional orientation with respect to replication. To control for gene expression levels, both the head-on and co-directional version of each gene was placed under the control of the same promoter (e.g. P_*spank(hy)*_).

In order to measure the relative association of type II topoisomerases with the conflict regions, we used a GFP fusion to the GyrA subunit of gyrase (Tadesse and Graumann 2006) and constructed a 3xMyc fusion to the ParC subunit of Topo IV. We expressed an IPTG-inducible *lacZ* gene in either the head-on or the co-directional orientation, and performed ChIP-Seq experiments in order to obtain a high resolution map of the association of type II topoisomerases with the engineered conflict regions. We found that both gyrase and Topo IV are preferentially enriched at the engineered conflict locus when the orientation of *lacZ* is head-on (Figure 1). Importantly, this enrichment was transcription-dependent. When we measured enrichment of these topoisomerases using ChIP-qPCR, we found that in the absence of the inducer, IPTG, the levels of topoisomerases at the engineered conflict regions were similar in the two orientations (Supplementary Figure 1). Furthermore, we confirmed that the GyrA signal was specific by performing control ChIPs of GFP only (unfused to GyrA) and found no enrichment at the *lacZ* gene in either orientation. It is noteworthy that we utilized standard formaldehyde crosslinking for the GyrA ChIPs. However, we were unable to ChIP ParC using formaldehyde. The ParC association was only detectable when we performed the ChIP assays using ciprofloxacin crosslinking, which specifically crosslinks active type II topoisomerases on DNA.

**Figure 1.**
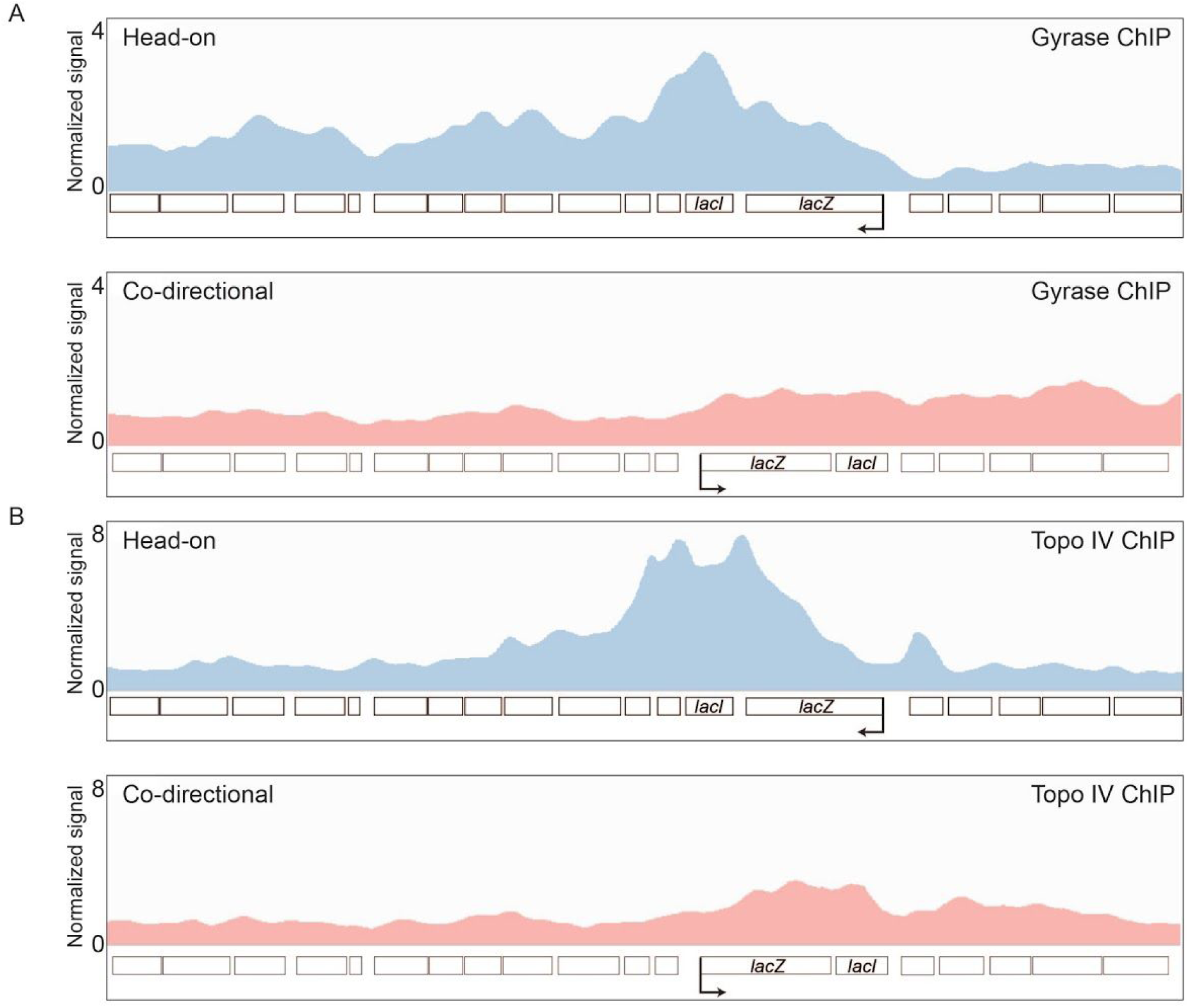
Type II topoisomerases are enriched at head-on genes. **(A)** DNA gyrase and **(B)** Topo IV ChIP-Seq profiles of cells carrying either a head-on (HO, blue, strain HM3863 (gyrase), HM4074 (ParC)) or co-directional (CD, red, strain HM3864 (gyrase), HM4075 (ParC)) *lacZ* engineered conflict. The direction of DNA replication is left to right. Direction of transcription is indicated by the promoter arrow on *lacZ*.

### Inhibition of type II topoisomerases increases replisome enrichment/replication stalling at head-on but not co-directional genes

In *E. coli*, gyrase and Topo IV promote replication fork progression (Khodursky et al. 2000). If torsional stress is a major problem at head-on conflict regions, then subtle inhibition of these topoisomerases should lead to increased replication fork stalling at head-on conflict regions. We tested this hypothesis by performing ChIP-seq of the replisome protein, DnaC, as a proxy for replication stalling. If fork progression is unimpeded, the distribution of DnaC enrichment should be equal along the genome in asynchronous bacterial cultures. We have demonstrated previously that DnaC enrichment is a good proxy for replication fork stalling (Lang et al. 2017; H. Merrikh et al. 2011; C. N. Merrikh, Brewer, and Merrikh 2015). To inhibit type II topoisomerase activity, we used subinhibitory doses of the antibiotic novobiocin. Novobiocin is a competitive inhibitor of type II topoisomerase ATPase activity (Sugino et al. 1978; Hardy and Cozzarelli 2003; Maxwell 1993). We performed ChIP-Seq experiments, where we measured the association of DnaC with the engineered conflict regions in media with and without sublethal concentrations of novobiocin (375 ng/mL). In untreated cells, we found preferential association of DnaC with the engineered conflict region in the head-on orientation (Figure 2, top panel). When the cells were treated with novobiocin, there was an increase in DnaC enrichment at the head-on but not the co-directional conflict region. These results suggest that without type II topoisomerase activity, topological problems at head-on genes can impede replication.

**Figure 2.**
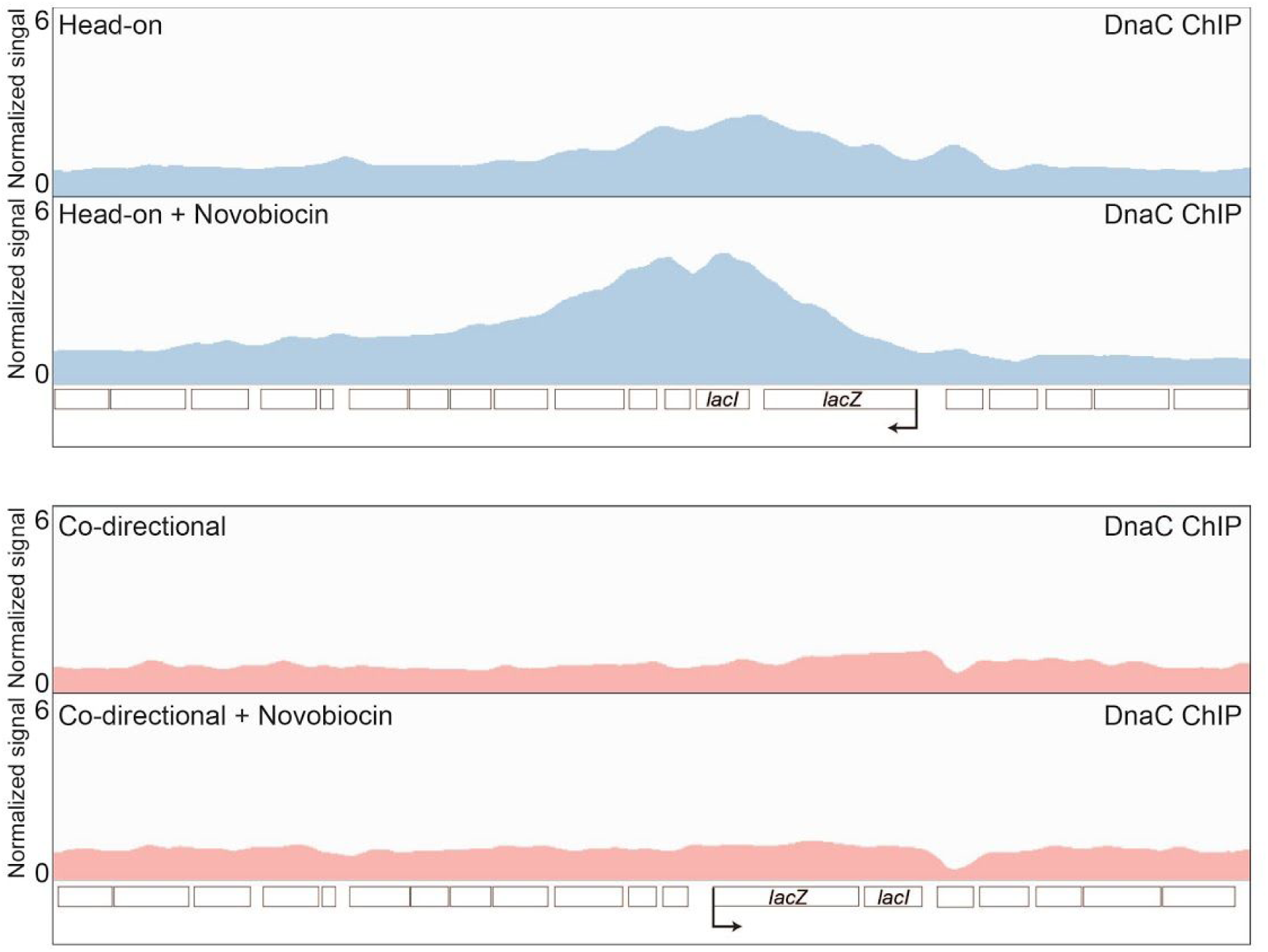
Type II topoisomerase inhibition results in increased stalling at head-on genes. **(A)** Representative DnaC ChIP-Seq profiles of cells carrying either a head-on (HO, blue, strain HM1300) or co-directional (CD, red, strain HM1416) *lacZ* engineered conflict, with and without novobiocin treatment (375 ng/mL). The direction of DNA replication is left to right. Direction of transcription is indicated by the promoter arrow on *lacZ*.

### Sublethal amounts of novobiocin compromises cell survival specifically in the presence of a strong head-on conflict

We previously showed that in the absence of critical conflict resolution factors, head-on conflicts can significantly compromise survival efficiency (Lang et al. 2017; Million-Weaver, Samadpour, and Merrikh 2015; C. N. Merrikh, Brewer, and Merrikh 2015). If type II topoisomerases are indeed important for conflict resolution, then the inhibition of these enzymes should impact survival of cells experiencing head-on conflicts. To test this hypothesis, we measured survival efficiency using colony forming units (CFUs) of cells containing the engineered conflicts, in the head-on or the co-directional orientation, upon chronic treatment with various concentrations of novobiocin. In the absence of novobiocin, there was no difference in survival efficiency of cells containing the engineered conflict in either orientation and regardless of whether the *lacZ* gene was transcribed (Figure 3A). When the cells were plated on novobiocin, again, there was no difference in survival efficiency between cells carrying the head-on or co-directional *lacZ* when transcription was off. However, when transcription was turned on, the cells carrying the head-on but not the co-directionally oriented *lacZ* gene were sensitized to low doses of novobiocin. The effects of head-on conflicts on survival, in response to inhibition of type II topoisomerases, was not specific to the chromosomal location or the nature of the gene used to induce the conflict.

**Figure 3.**
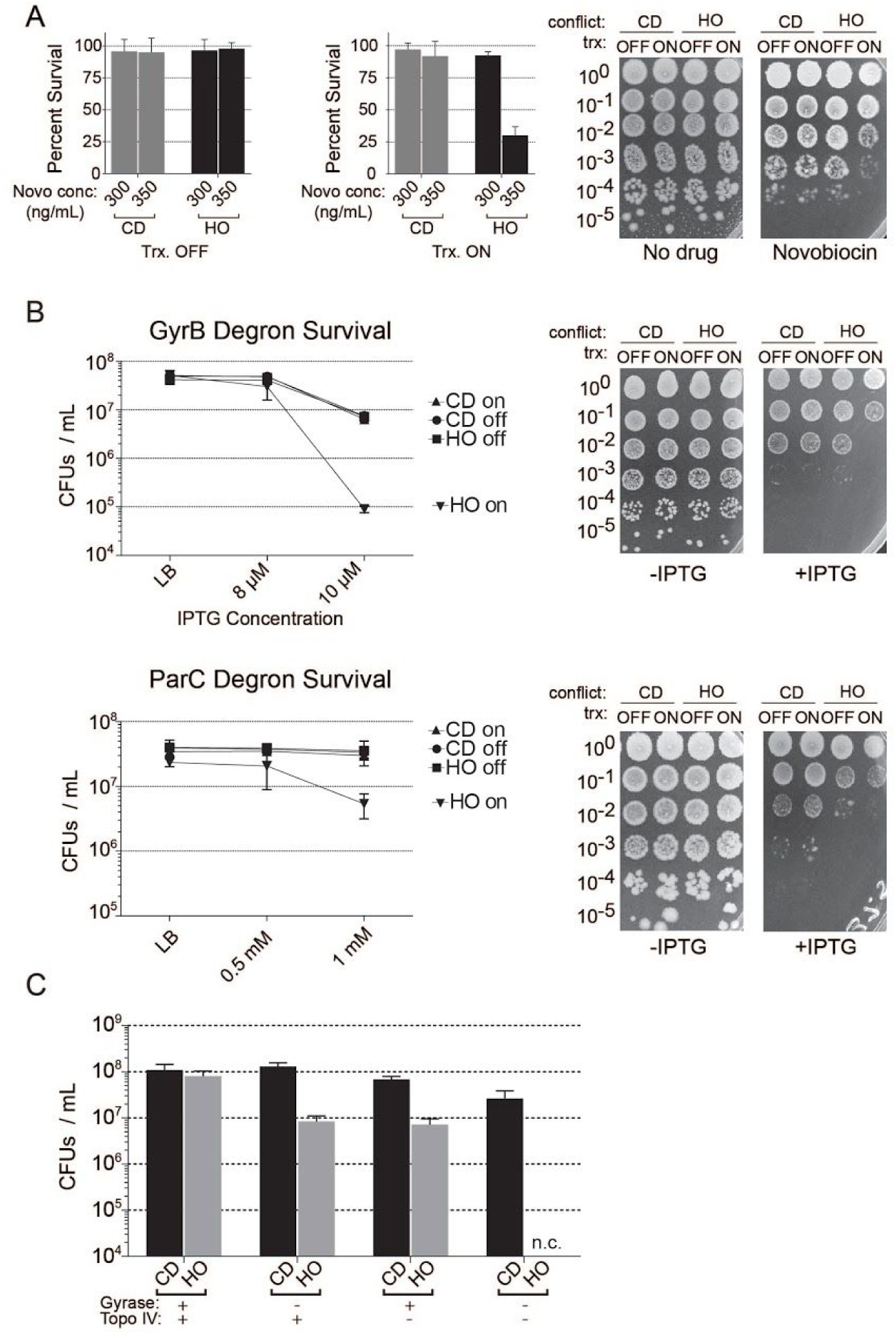
DNA gyrase and Topo IV act in parallel to resolve head-on conflicts. **(A)** Survival of cells harboring either a repressed (head-on, HO, HM640 or co-directional, CD, HM1794) or constitutively transcribed (head-on, HO, HM211 or co-directional, CD, HM1795) *lacZ* engineered conflict plated on LB or LB supplemented with novobiocin (375 ng/mL). Bar graphs are quantification (mean and standard deviation) of three independent biological replicates. Survival after conditional (IPTG dependant) depletion of either **(B)** gyrase or **(C)** Topo IV in cells harboring either a repressed (head-on, HO, HM1951/HM1467 or co-directional, CD, HM1949/HM1468) or constitutively transcribed (head-on, HO, HM1952/HM1450 or co-directional, CD, HM1950/HM1469) *lacZ* engineered conflict plated on LB or LB supplemented with IPTG (as indicated). **(D)** Survival of cells harboring a novobiocin resistant *gyrB* allele, a conditional gyrase depletion (IPTG dependent) system and a constitutively transcribed (head-on, HO, HM2420 or co-directional, CD, HM2421) *lacZ* engineered conflict plated on LB, LB supplemented with novobiocin (7 µg/mL), LB supplemented with IPTG (10 µM), or both novobiocin and IPTG.

In order to control for potential indirect effects of genomic context, chromosomal location, and sequence, we performed similar survival experiments using a second engineered conflict system. In this system, we inserted a different transcription unit, the *luxABCDE* operon, onto the opposite (right) arm of the chromosome. We performed the survival experiments with this system as described above. The results of these experiments were consistent with the *lacZ* system: there was a survival defect in cells containing the *luxABCDE* operon, but only when this transcription unit was in the head-on orientation, and only when the genes were transcribed (Supplementary Figure 2).

### Both gyrase and Topo IV are critical for the resolution of head-on conflicts

Novobiocin has activity against both gyrase and Topo IV, although the affinity of the drug for Topo IV is much weaker than that for gyrase (Peng and Marians 1993; Sugino et al. 1978). It was unclear from our survival assays whether the survival defects were a result of inhibition of only gyrase, or TopoIV, or both. It is likely that only gyrase activity is inhibited at the concentrations of novobiocin we used in our experiments. However, it can’t be ruled out that Topo IV activity is also inhibited to some extent under these conditions. To directly determine the contribution of each of the two enzymes to conflict resolution, we adapted a conditional degradation system (Griffith and Grossman 2008) to specifically deplete the GyrB subunit of gyrase or the ParC subunit of Topo IV. This system is induced by IPTG. In order to detect potentially subtle differences in survival of our engineered conflict strains, we used concentrations of IPTG that only slightly depleted GyrB, and subtly impacted survival of wild-type cells (gyrase is essential, so a complete depletion cannot be used here). We then tested the survival of cells carrying engineered conflicts under these conditions, but now the engineered conflicts expressed *lacZ* from a different promoter, P_*xis*_, which is constitutively active. The “transcription off” control for this engineered conflict is achieved through the use of a strain where this promoter is constitutively off. In both the GyrB and ParC degron systems, we found that without IPTG, there was no difference in survival efficiency, in any of the engineered conflict strains. When we specifically depleted GyrB in cells carrying the co-directional conflict, transcription of *lacZ* made no difference in survival efficiency. In cells carrying the head-on reporter, however, there was about a 2-log defect in survival, only when the transcription of the reporter was on (Figure 3B). Similarly, when we depleted ParC, there was about a 90% reduction in the number of CFUs when comparing strains with transcriptionally active versus inactive head-on *lacZ*.

In order to address whether gyrase and Topo IV act together or in parallel, we constructed a strain that had a mutation in the *gyrB* gene that conferred high level of resistance to novobiocin (R128L). In this background, novobiocin treatment can only impact Topo IV. In this same strain, we fused the *gyrB* gene to the *ssrA* tag in order to deplete gyrase with our degron system. We found that low concentrations of IPTG (GyrB depletion) or high levels of novobiocin (Topo IV inhibition) both led to a survival defect in the strain carrying the head-on but not the co-directional conflict (Figure 3C). When we treated cells with both IPTG and novobiocin, the cells expressing the head-on *lacZ* gene were not viable (Figure 3C). This result indicates that gyrase and Topo IV are the only two factors that can resolve the torsional stress problem at head-on conflict regions.

### Inhibition of type II topoisomerases reduces R-Loop formation at head-on conflict regions

There is evidence in the literature that topoisomerase activity can influence R-Loop formation, at least *in vitro* and in human cells (Massé and Drolet 1999; Tuduri et al. 2009). Furthermore, our results described above strongly suggest that DNA topology is a serious problem at head-on conflict regions. Given our prior results that R-Loops contribute to many of the detrimental outcomes of head-on conflicts, we decided to investigate whether resolution of head-on conflicts by topoisomerases influence R-Loop formation. We tested this hypothesis by directly measuring R-Loop levels at the conflict regions, in strains lacking RNase HIII (Lang et al. 2017; Naoto Ohtani et al. 1998; Randall, Hirst, and Simmons 2018). We performed DNA-RNA Hybrid ImmunoPrecipitations coupled to deep sequencing (DRIP-Seq) experiments using the S9.6 antibody, which recognizes RNA:DNA hybrids. Consistent with what we have measured previously using qPCR (Lang et al. 2017), we found more R-loops when the *lacZ* gene was expressed in the head-on orientation compared to the co-directional orientation (Figure 4A). We then used DRIP-Seq to measure R-loops in cells treated with low levels of novobiocin to subtly reduce the activities of both type II topoisomerases. Remarkably, we found that when type II topoisomerases are inhibited, R-loop levels are reduced at head-on conflict regions. As a control, we looked at expression levels of the reporter gene using ChIP-Seq of the beta subunit of RNAP, RpoB. We found no difference in RpoB occupancy at the engineered conflict regions with novobiocin treatment, indicating that the lowered R-loop levels are not simply due to reduced expression of the head-on *lacZ* gene (Supplementary Figure 3).

**Figure 4.**
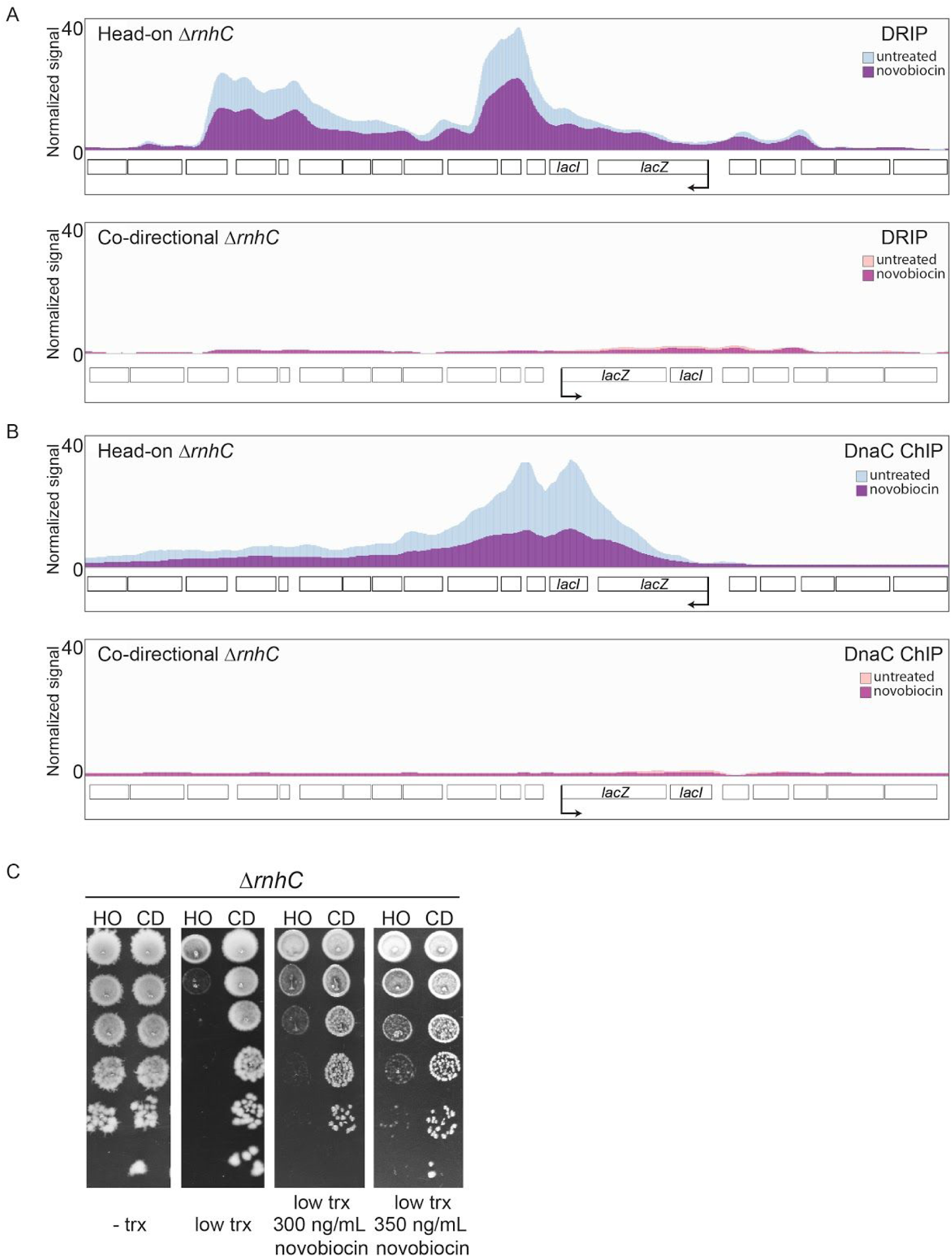
Resolution of head-on conflicts by type II topoisomerases drives the formation of toxic R-loops. **(A)** DRIP-Seq and **(B)** DnaC ChIP-Seq profiles of cells lacking RNase HIII harboring either a head-on (HO, blue, strain HM2043) or co-directional (CD, red, strain HM2044) *lacZ* reporter treated or untreated with novobiocin. **(C)** Survival of cells lacking RNase HIII harboring either a head-on (HO, blue, strain HM2043) or co-directional (CD, red, strain HM2044) *lacZ* reporter treated or untreated with novobiocin.

### Inhibiting type II topoisomerases rescues R-Loop mediated replisome stalling

R-loops at head-on genes stall the replisome in many different organisms (Prado and Aguilera 2005; Lang et al. 2017; Hamperl et al. 2017). If type II topoisomerase activity is driving R-loop formation at head-on genes, then treating cells with low doses of novobiocin should reduce replisome stalling at head-on conflict regions in cells lacking RNase HIII. We tested this hypothesis using DnaC ChIP-Seq, as described above. As we published previously, we found that there is a preferential association of DnaC with head-on versus co-directional conflict regions, and this difference is significantly increased in cells lacking RNase HIII (Figure 4B). This DnaC ChIP signal at head-on conflict regions, in cells lacking RNase HIII, corresponds to complete replication fork stalling at that locus (Lang et al. 2017). When we treated cells with low amounts of novobiocin to inhibit topoisomerase activity, there was a marked decrease in DnaC enrichment at the head-on conflict region (Figure 4B). This result suggests that the type II topoisomerases are responsible for R-Loop mediated replisome stalling at head-on conflict regions.

### Inhibiting type II topoisomerases rescues death by R-Loops

We previously showed that increased stalling due to unresolved R-loops at head-on genes is lethal (Lang et al. 2017). If topoisomerase activity is driving R-loop formation at head-on genes, then limiting that activity should increase the viability of cells that contain an engineered head-on conflict and lack RNase HIII. We tested this model by measuring the viability of cells lacking RNase HIII, and expressing either the head-on or co-directional *lacZ* in the presence of low concentrations of novobiocin. As expected, cells with the co-directional reporter had no growth defect when the *lacZ* gene was induced with IPTG. In contrast, cells expressing the *lacZ* gene in the head-on orientation had significant cell survival defects. Remarkably, chronic novobiocin exposure rescued these defects in a dose dependent manner (Figure 4C). Altogether, these results suggest that the resolution of head-on conflicts by type II topoisomerase activity is driving toxic R-loop formation.

### Introduction of negative supercoils by gyrase promotes toxic R-Loop formation at head-on conflict regions

Novobiocin inhibits both gyrase and Topo IV activity. Because gyrase is much more sensitive to novobiocin than Topo IV, we wondered whether the decreased R-loop levels was due to inhibition of gyrase, and not inhibition of Topo IV or pleiotropic effects of novobiocin. Gyrase has two activities: 1) relaxation of positive supercoiling and 2) introduction of negative supercoiling (Vos et al. 2011). Both *in vitro* and *in vivo*, R-loops have been shown to form more readily (or are more stable) in the presence of gyrase (Massé and Drolet 1999; Drolet, Bi, and Liu 1994; Drolet et al. 1995). This is likely due to the introduction of negative supercoiling by gyrase, as negatively supercoiled DNA will energetically favor R-loop formation, although recent work has suggested that highly positively increased supercoiling could also impact R-Loop formation (Stolz et al. 2019). We tested this model by utilizing the *gyrB* (R138L) mutant, which has reduced ATPase activity and thus has a ten-fold reduction in the ability to introduce negative supercoils (Contreras and Maxwell 1992; Gross et al. 2003). Whether and/or how much this mutation impacts the positive supercoil relaxation activity of gyrase has not been assessed. However, Topo IV can resolve torsional stress at conflict regions in parallel to gyrase as we showed above. Therefore, even if the positive supercoil relaxation activity of gyrase is impacted by the R138L mutation, the major effect of this mutation at the conflict region will be a loss of negative supercoil introduction. We used survival assays to measure viability of Δ*rnhC* strains containing the mutant *gyrB*, in the presence of either the head-on or co-directionally oriented conflicts. As expected, there was no effect of transcription on the viability of the cells carrying the co-directional reporter construct. Consistent with our previous work, we found that induction of the conflict reporter was completely lethal when it was oriented head-on to replication. Remarkably, we found that the *gyrB* R138L mutation completely rescued this lethality (Figure 5A). We tested whether this rescue was due to the reduction of stable R-loops at the conflict region by measuring R-loops directly by DRIP-qPCR experiments. In cells lacking RNase HIII, consistent with what we have previously reported, we measured about a 5-fold higher R-loop at the head-on compared to the co-directional *lacZ* (Figure 5B). When we measured R-loops in cells with the R138L *gyrB* mutation, the R-loop levels were similar at the head-on and co-directional conflict regions. These results demonstrate that it is specifically the introduction of negative supercoils by gyrase at head-on conflict regions that leads to the formation (and/or stability) of toxic R-loops.

**Figure 5.**
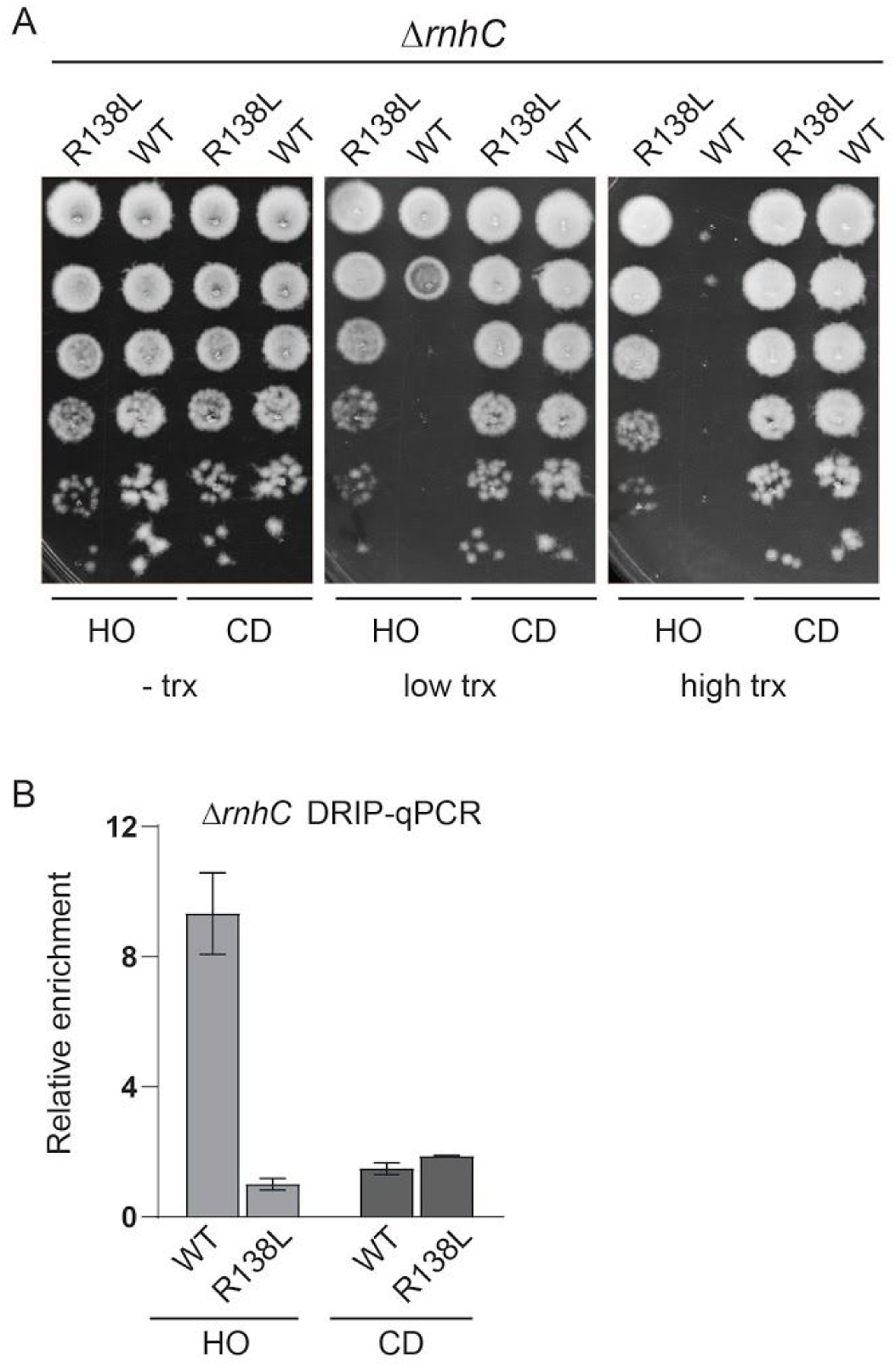
The negative supercoiling activity of DNA gyrase results in R-loop formation at head-on genes. **(A)** Survival of cells lacking RNase HIII with either the WT or R138L *gyrB* allele harboring either a head-on (HO, HM2043/HM4065) or co-directional (CD, HM2044/HM4066) *lacZ* engineered conflict. **(B)** DRIP-qPCR analysis of cells lacking RNase HIII with either the WT or R138L *gyrB* allele harboring either a head-on (HO, HM2043/HM4065) or co-directional (CD, HM2044/HM4066) *lacZ* reporter.

## Discussion

Our results strongly suggest that positive supercoils build up at head-on conflict regions. We find that type II topoisomerases are enriched at head-on genes. Furthermore, their depletion results in gene orientation-specific and transcription-dependent replisome stalling and cell viability defects in cells with highly expressed head-on genes. These viability defects likely stem from increased stalling of the replication fork that we observe at head-on genes in cells where type II topoisomerases are inhibited. However, we also find that gyrase activity at head-on genes drives R-loop formation.

Our results can be explained by a “spin-diffusion” model, where excess negative supercoils generated by gyrase promote R-Loop formation through the diffusion of the supercoils past RNAPs (Figure 6). This process is initially triggered by positive supercoil buildup between the replication and transcription machineries at head-on conflict regions, which is rapidly removed by type II topoisomerases. Gyrase would convert the conflict region to hyper-negatively supercoiled DNA (Lynch and Wang 1993; Drolet, Bi, and Liu 1994; Drolet et al. 1995). This increase in negative supercoiling could diffuse through RNAP spinning about its axis (Nudler 2009, 2012). Diffusion of the negative supercoils will increase the chance for R-loop formation behind RNAP. R-loops would then have to be processed by RNase H enzymes. Alternatively, the sudden release of torsional strain by type II topoisomerases could cause RNAP to rapidly progress, generating excessive negative supercoils and R-loop formation (Kuzminov 2017). The importance of Topo IV adds a second dimension to our model. The observations that Topo IV is important for conflict resolution can be interpreted in two ways: 1) Topo IV helps relaxes positive supercoils at conflict regions, or 2) the increased torsional stress leads to the formation of catenanes by inducing replisome spinning about its axis. Given that there is a significant amount of literature showing that Topo IV is critical for catenane resolution, we favor the second possibility (Figure 6). These models however are not mutually exclusive and it is still very much possible that Topo IV also relaxes positive supercoils at head-on conflict regions.

**Figure 6.**
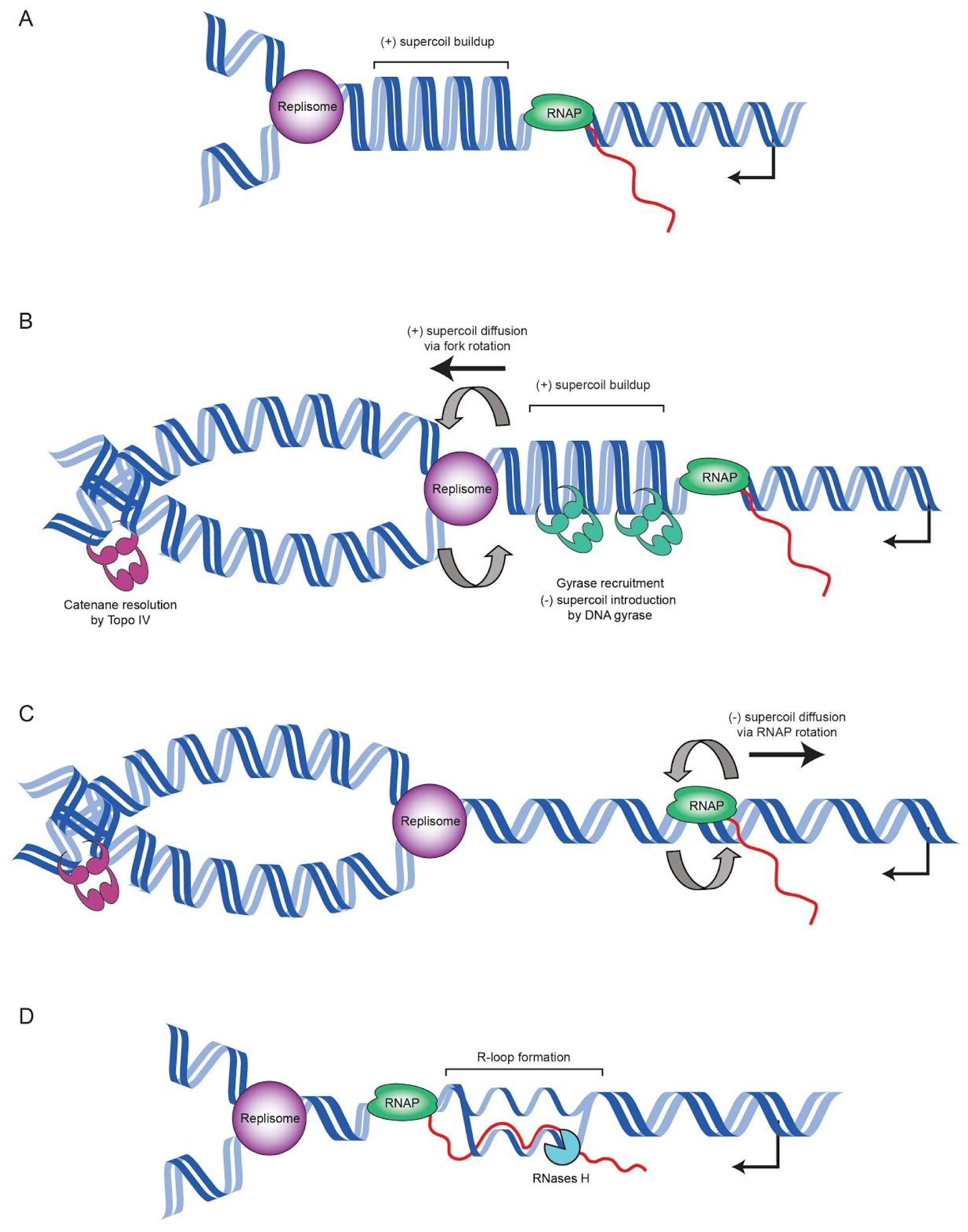
Proposed model for topological changes and R-loop formation at head-on conflict regions. **(A)** As the replisome and head-on transcription unit converge, positive supercoils accumulate in between the two machineries. **(B)** DNA gyrase resolves the positive supercoil buildup. The replisome also likely spins to relieve the torsional strain, producing catenanes behind the replication fork, which are resolved by Topo IV. **(C)** Gyrase activity rapidly converts the conflict region to negatively supercoiled DNA causing RNAP to spin about its axis. Negative supercoils diffuse behind RNAP. **(D)** The diffused negative supercoils drive R-loop formation behind RNAP, which are resolved by RNAse H enzymes.

The problem of replication-transcription conflicts exists in all domains of life. Gene-orientation dependent effects of transcription on DNA replication have been a topic of interest since Sarah French’s discovery in 1992 that a head-on gene slows replication significantly more than a co-directional gene (French 1992). However, why the orientation of transcription relative to DNA replication matters has remained a mystery. The protein makeup of the two machineries are the same in both orientations, yet the direction in which they encounter each other has profound downstream effects. One potential hypothesis that could explain gene orientation effects of conflicts has to do with the strand specificity of where the replicative helicase resides (lagging strand in bacteria, leading strand in eukaryotes) (Hamperl and Cimprich 2016; Gómez-González and Aguilera 2019). This model could explain why the two different types of conflicts between the replisome and RNA polymerases have differential consequences. However, the discovery that R-loops are a major problem in head-on but not co-directional conflicts in both bacteria and mammalian cells undermines this model (Lang et al. 2017; Hamperl et al. 2017). The replicative helicase moves on the lagging strand in bacteria whereas it moves on the leading strand in mammalian cells. Yet, the fundamental problem of R-Loop enrichment in head-on conflicts remains the same across these species. Therefore, gene-orientation specific problems are unlikely to stem from this particular architectural feature of the replisome complex. On the other hand, production of positive supercoils by the replication and transcription machineries is a universal feature, and therefore, could be the fundamental mechanism underlying gene orientation-specific effects of replication-transcription conflicts.

It is clear from this work, as well as others’, that after an encounter with the transcription machinery, replication stalls, the replisome collapses, and replication progression requires restart proteins (Mangiameli et al. 2017). However, the extent to which the fork is remodeled and whether there is replication fork reversal after a head-on conflict is not yet clear. Previous studies have implied that in head-on conflicts, the replication fork reverses, and is subsequently processed by recombination proteins (Million-Weaver, Samadpour, and Merrikh 2015; De Septenville et al. 2012). Furthermore, it has been shown *in vitro* that replication forks reverse in response to positive supercoil accumulation (Postow, Ullsperger, et al. 2001). Given that at least in eukaryotic systems supercoiling can push the fork back, our data presented here is consistent with the model that conflicts lead to replication fork reversal due to positive supercoil buildup.

We previously showed that R-loops are a major problem for replication forks that are approaching an actively transcribed head-on gene (Lang et al. 2017; Hamperl et al. 2017). Here, we find that R-loop formation and/or stabilization at head-on genes stems from the introduction of negative supercoils by gyrase at these regions. This phenomenon has important implications. Most importantly, our previous work demonstrated that the increased mutagenesis of head-on genes is driven by R-Loops in wild-type cells. Given that gyrase activity is facilitating R-loop formation, our results suggest that the activity of this enzyme leads to increased mutagenesis, albeit indirectly. Interestingly, as our group and others have shown, the full capacity of gyrase to introduce negative supercoils is not essential for viability (Gross et al. 2003). Why then, is this function conserved? We previously proposed that the head-on orientation is retained for some genes as a mechanism to increase mutagenesis and promote gene specific evolution (Paul et al. 2013; C. N. Merrikh and Merrikh 2018). We speculate that the introduction of negative supercoils by gyrase is a highly conserved function across bacteria at least partially because it is evolutionarily beneficial. In particular, we showed previously that head-on genes, including many of the critical stress response genes, evolve faster than co-directional genes. Under selection, these head-on genes will likely gain beneficial mutations faster than if they were co-directionally oriented, simply due to the increased mutation rates which are facilitated by conflicts. If those beneficial mutations are obtained through negative supercoil introduction by gyrase (and downstream R-Loop formation), this property of gyrase would be retained over evolutionary time despite the fact that it is not immediately necessary for viability. In other words, this activity of gyrase would hitchhike along in cells that have rapidly adapted to their environment by obtaining beneficial mutations relatively quickly.

In this work, we discovered (what appears to be) the main source of gene orientation-specific problems in replication-transcription conflicts. We also unraveled an intriguing feature of topoisomerases that in the big picture, could place them into a category of evolutionarily beneficial factors that increase mutagenesis. These findings highlight the fundamental importance and influence of conflicts and DNA supercoiling on cellular physiology, genome organization, and adaptation.

## Author contributions

K.S.L. and H.M. designed and performed the experiments, analyzed the data, and wrote the paper.

## Acknowledgements

We would like to thank Christopher Merrikh and the other members (past and present) of the Merrikh lab, as well as Neil Osheroff and Felipe Cortés Ledesma for helpful discussions. We would also like to thank Peter Graumann for sharing strains. This work was supported by the NIGMS DP2GM110773 award from the National Institutes of Health and the Research Royalty Funds from the University of Washington to H.M., and the 5T32-AI055396-13 Bacterial Pathogenesis Training Grant (University of Washington) award and the 1F32 AI140557-01 Ruth L. Kirschstein National Research Postdoctoral award to K.S.L.. We would like to dedicate this paper to the late James Champoux, who discovered Top1, and discussed this project with us on various occasions during our time at the University of Washington.

## Figure and table legends

**Supplementary Figure 1.**
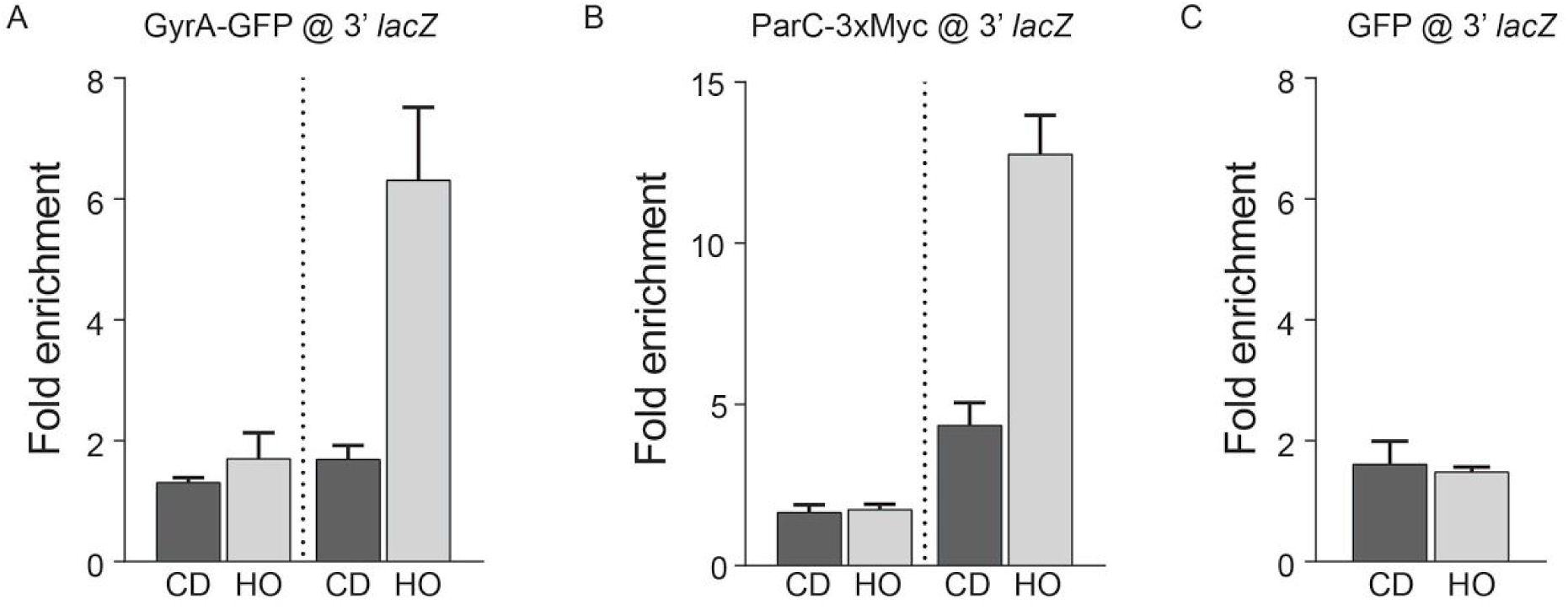
Association of type II topoisomerases with head-on genes is dependant on transcription. ChIP-qPCR analysis of cells expressing either the head-on or co-directional engineered conflict. Transcription of the engineered conflict was either uninduced or induced with IPTG. Enrichment of either **(A)** gyrase or **(B)** Topo IV was measured using qPCR targeting the engineered conflict gene *lacZ* compared to a control locus *yhaX*. **(C)** To control for the use of GFP as a fusion tag, GFP alone was expressed in cells expressing the engineered conflicts and its enrichment was measured by ChIP-qPCR.

**Supplementary Figure 2.**
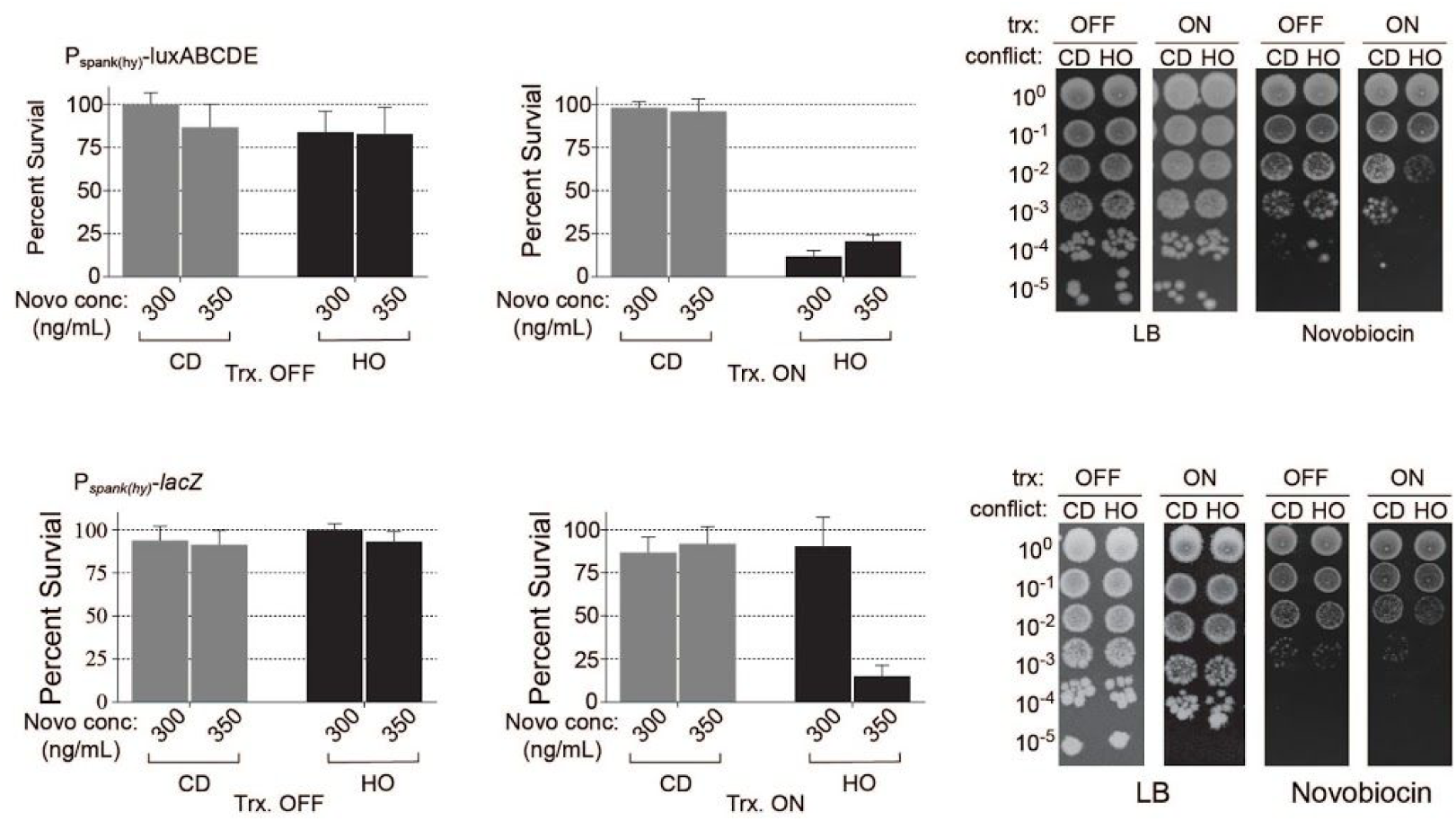
Sensitivity of cells expressing a highly transcribed gene is independent of gene sequence and genomic location. Novobiocin survival assays of cells expressing P_*spank(hy)*_*-luxABCDE* and P_*spank(hy)*_*-lacZ* at the *amyE* locus. Quantification is shown as percent survival. Representative plates of the highest novobiocin concentration (350 ng/mL) are shown.

**Supplementary Figure 3.**
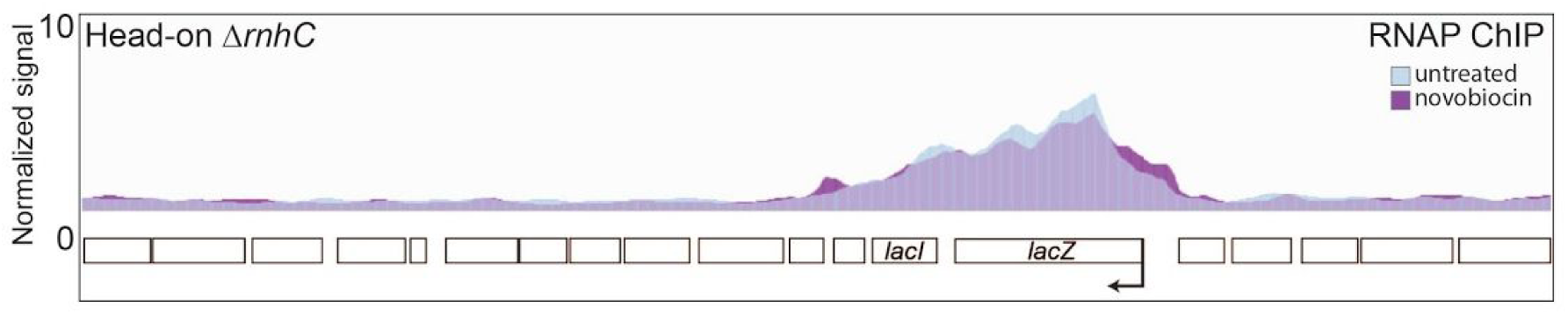
RNAP occupancy at head-on genes doesn’t change with treatment of low doses of novobiocin. Representative RNAP ChIP-seq plot of Δ*rnhC* cells expressing the head-on engineered conflict untreated (blue) and treated with novobiocin (magenta).

**Table 1.** Strains, plasmids, and primers used in this study.

| Strain | Genotype | Reference |
| --- | --- | --- |
| HM1 | phe trp | Brehm et al 1973 |
| HM211 | phe trp *thrC*::P<sub>xis</sub>-*lacZ* (HO) *ICEBs1*(0) | Merrikh et al 2015 |
| HM640 | phe trp *thrC*::P<sub>xis</sub>-*lacZ* (HO) | Merrikh et al 2015 |
| HM1300 | phe trp *amyE*::P<sub>spank(hy)</sub>-*lacZ* (HO) | Lang et al 2017 |
| HM1416 | phe trp *amyE*::P<sub>spank(hy)</sub>-*lacZ* (CD) | Lang et al 2017 |
| HM1450 | phe trp *thrC*::P<sub>xis</sub>-*lacZ* (HO) *ICEBs1*(0) *amyE*::P<sub>spank(hy)</sub>-*sspB*<br>*parC*::*parC-ssrA* | This study |
| HM1467 | phe trp *thrC*::P<sub>xis</sub>-*lacZ* (HO) *amyE*::P<sub>spank(hy)</sub>-*sspB* *parC*::*parC-ssrA* | This study |
| HM1468 | phe trp *thrC*::P<sub>xis</sub>-*lacZ* (CD) *amyE*::P<sub>spank(hy)</sub>-*sspB* *parC*::*parC-ssrA* | This study |
| HM1469 | phe trp *thrC*::P<sub>xis</sub>-*lacZ* (CD) *ICEBs1*(0) *amyE*::P<sub>spank(hy)</sub>-*sspB*<br>*parC*::*parC-ssrA* | This study |
| HM1949 | phe trp *thrC*::P<sub>xis</sub>-*lacZ* (CD) *amyE*::P<sub>spank(hy)</sub>-*sspB* *gyrB*::*gyrB-ssrA* | This study |
| HM1950 | phe trp *thrC*::P<sub>xis</sub>-*lacZ* (CD) *ICEBs1*(0) *amyE*::P<sub>spank(hy)</sub>-*sspB*<br>*gyrB*::*gyrB-ssrA* | This study |
| HM1951 | phe trp *thrC*::P<sub>xis</sub>-*lacZ* (HO) *amyE*::P<sub>spank(hy)</sub>-*sspB* *gyrB*::*gyrB-ssrA* | This study |
| HM1952 | phe trp *thrC*::P<sub>xis</sub>-*lacZ* (HO) *ICEBs1*(0) *amyE*::P<sub>spank(hy)</sub>-*sspB*<br>*gyrB*::*gyrB-ssrA* | This study |
| HM2043 | phe trp *amyE*::P<sub>spank(hy)</sub>-*lacZ* (HO) *ΔrnhC*::*MLS* | Lang et al 2017 |
| HM2044 | phe trp *amyE*::P<sub>spank(hy)</sub>-*lacZ* (CD) *ΔrnhC*::*MLS* | Lang et al 2017 |
| HM2420 | phe trp *thrC*::P<sub>xis</sub>-*lacZ* (HO) *ICEBs1*(0) *amyE*::P<sub>spank(hy)</sub>-*sspB*<br>*gyrB*::*gyrB(R138L)-ssrA* | This study |
| HM2421 | phe trp *thrC*::P<sub>xis</sub>-*lacZ* (CD) *ICEBs1*(0) *amyE*::P<sub>spank(hy)</sub>-*sspB*<br>*gyrB*::*gyrB(R138L)-ssrA* | This study |
| HM2655 | phe trp *ΔrnhC*::*MLS* | Lang et al 2017 |
| HM3863 | phe trp *amyE*::P<sub>spank(hy)</sub>-*lacZ* (HO) *gyrA*::*gyrA-gfp* | This work |
| HM3864 | phe trp *amyE*::P<sub>spank(hy)</sub>-*lacZ* (CD) *gyrA*::*gyrA-gfp* |  |
| HM3387 | *phe trp gyrB(R138L)* | Samadpour et al |
|  |  | 2018 |
| HM4064 | phe trp $\Delta rnhC::MLS$ gyrB(R138L) | This study |
| HM4065 | phe trp $\Delta rnhC::MLS$ gyrB(R138L) amyE::P<sub>spank(hy)</sub>-lacZ (HO) | This study |
| HM4066 | phe trp $\Delta rnhC::MLS$ gyrB(R138L) amyE::P<sub>spank(hy)</sub>-lacZ (CD) | This study |
| HM4074 | phe trp amyE::P<sub>spank(hy)</sub>-lacZ (HO) parC::parC-3xMyc | This study |
| HM4075 | phe trp amyE::P<sub>spank(hy)</sub>-lacZ (CD) parC::parC-3xMyc | This study |

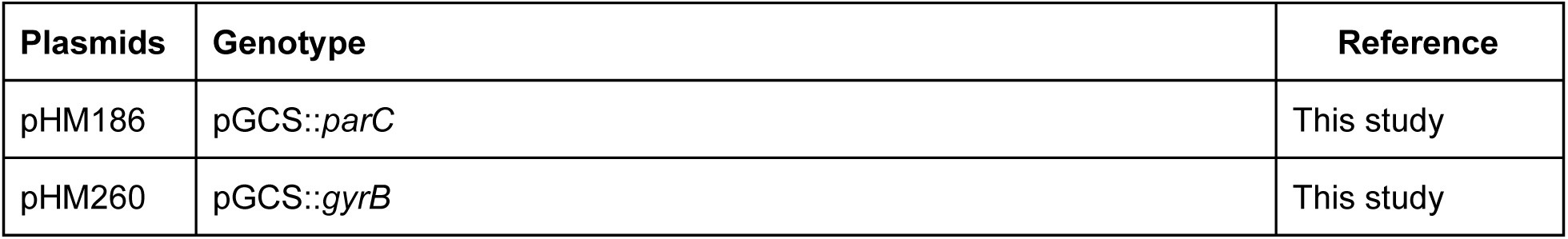
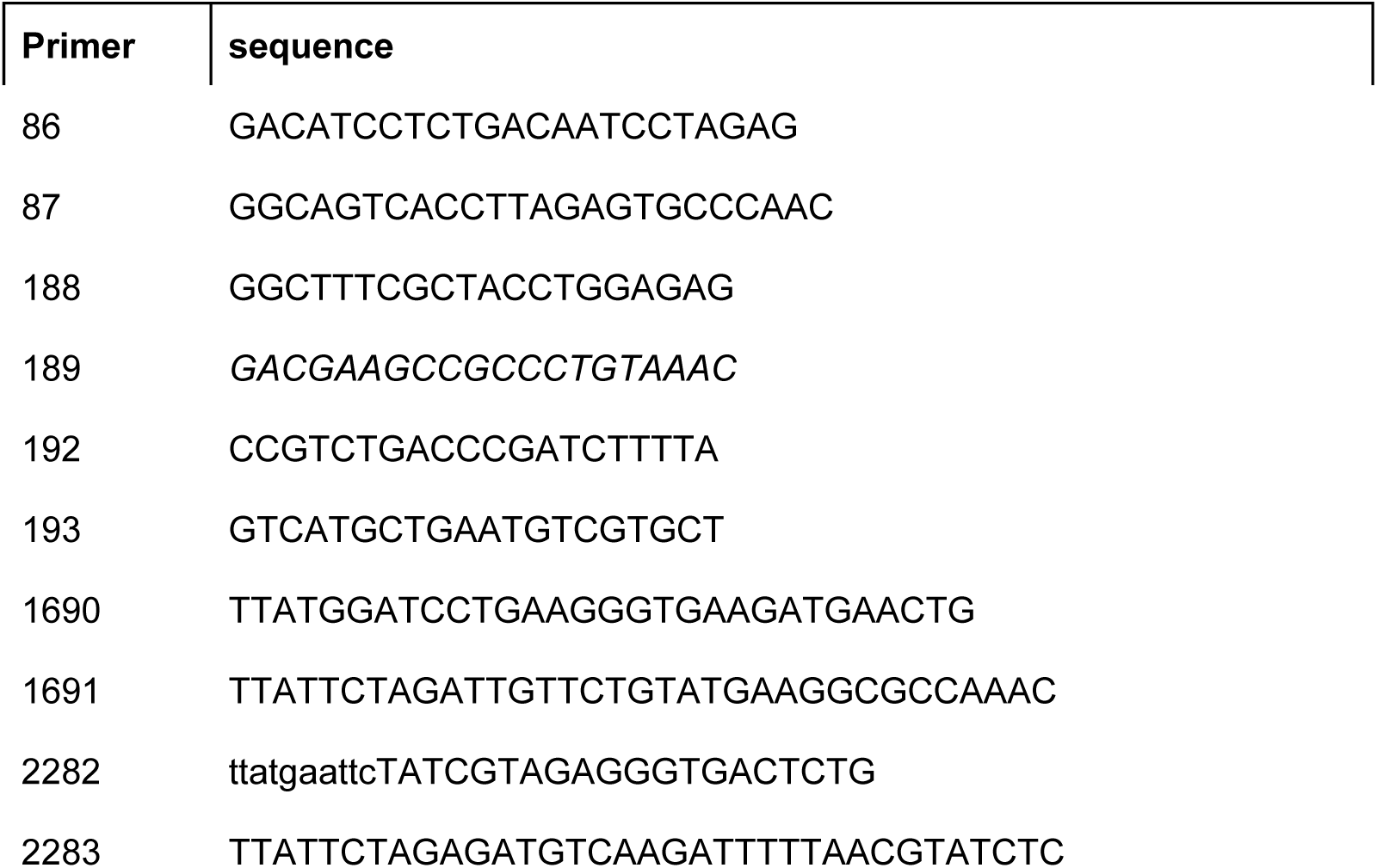

## Materials and methods

### Bacterial strains and growth conditions

All strains were constructed in the HM1 (JH642) (Brehm et al., 1973) *B. subtilis* background. The *rnhC::mls* mutant (HM711) was obtained from the *Bacillus* genetic stock center (Columbus, OH). To move the *rnhC::mls* allele, genomic DNA was extracted from HM711 using a commercially available kit (Thermo) and used to transform into HM1 (and its derivatives with reporter constructs) as per standard protocol (Cutting and Harwood, 1990). Strains were streaked on LB agar plates and supplemented with antibiotics where appropriate. Precultures were inoculated from single colonies into 2 or 5 mL of LB broth and incubated at 37° C with shaking (260 RPM). Precultures were used to inoculate experimental cultures which were grown and treated as indicated for each different experiment in the materials and methods.

*E. coli* DH5α was used to propagate recombinant DNA vectors. Transformations were done using heat shock of competent *E. coli. E. coli* cultures were grown at 37° C with shaking (260 RPM) in LB supplemented with 50 µg/mL carbenicillin where appropriate. All plasmid vectors were purified using a commercially available plasmid extraction kit (Thermo).

### Plasmid and strain constructions

**pHM186** PCR was used to amplify 500 bp of the 3’ end of *parC* without the stop codon (primers HM1690/1691). The resulting amplicon was digested with BamHI and XbaI and ligated into pGCS (Griffith and Grossman 2008).

**pHM260** PCR was used to amplify 500 bp of the 3’ end of *gyrB* without the stop codon (primers HM22832284). The resulting amplicon was digested with EcoRI and XbaI and ligated into pGCS.

**HM1450** Strain HM867 (C. N. Merrikh, Brewer, and Merrikh 2015) was transformed with plasmid pHM186 and transformants were selected on LB plates containing chloramphenicol.

**HM1467** Strain HM866 (C. N. Merrikh, Brewer, and Merrikh 2015) was transformed with plasmid pHM186 and transformants were selected on LB plates containing chloramphenicol.

**HM1468** Strain HM868 (C. N. Merrikh, Brewer, and Merrikh 2015) was transformed with plasmid pHM186 and transformants were selected on LB plates containing chloramphenicol.

**HM1469** Strain HM869 (C. N. Merrikh, Brewer, and Merrikh 2015) was transformed with plasmid pHM186 and transformants were selected on LB plates containing chloramphenicol.

**HM1949** Strain HM868 was transformed with plasmid pHM190 and transformants were selected on LB plates containing chloramphenicol.

**HM1950** Strain HM869 was transformed with plasmid pHM190 and transformants were selected on LB plates containing chloramphenicol.

**HM1951** Strain HM866 was transformed with plasmid pHM190 and transformants were selected on LB plates containing chloramphenicol.

**HM1952** Strain HM867 was transformed with plasmid pHM190 and transformants were selected on LB plates containing chloramphenicol.

**HM2420** Strain HM866 was transformed with genomic DNA purified from HM3387 and transformants were selected on LB plates containing novobiocin (4 µg/mL). The novobiocin resistant transformant was then transformed with pHM260.

**HM2421** Strain HM869 was transformed with genomic DNA purified from HM3387 and transformants were selected on LB plates containing novobiocin (4 µg/mL). The novobiocin resistant transformant was then transformed with pHM260.

**HM4064** Strain HM3387 was transformed with gDNA purified from strain HM2655 and transformants were selected for on LB containing erythromycin and lincomycin.

**HM4065** Strain HM4064 was transformed with plasmid pHM171 (Lang et al. 2017).

**HM4066** Strain HM4064 was transformed with plasmid pHM180 (Lang et al. 2017).

### Viability assays - chronic treatments

Strains were struck on LB plates supplemented with the appropriate antibiotic from freezer stocks and incubated overnight at 37° C. Single colonies were used to inoculate 2 mL LB cultures in glass tubes. The cultures were grown at 37° C with shaking (260 RPM) to OD600 = 0.5-1.0. Precultures were adjusted to OD 0.3 and then serially diluted in 1x Spizzen’s Salts (15 mM ammonium sulfate, 80 mM dibasic potassium phosphate, 44 mM monobasic potassium phosphate, 3.4 mM trisodium citrate, and 0.8 mM magnesium sulfate). 5ul of each dilution was plated onto LB plates and incubated at 30° C overnight. For survival assays with reporter strains, LB plates were either supplemented or not with various concentrations of novobiocin and/or IPTG as indicated in the figure legends. For the type II topoisomerase degron experiments, chloramphenicol was added to the all of the media to maintain the stability of degron tag. Plates were imaged with a BioRad Gel Doc™ XR+ Molecular Imager^®^ and colonies were enumerated.

### Chromatin immunoprecipitation assays (ChIPs)

Precultures were diluted to OD600 of 0.05 in LB and grown at 30° C with shaking. At OD600 ∼0.1, cultures were induced with 1 mM IPTG (final concentration) and grown until the culture was at OD600 = 0.3 and processed as described (Merrikh et al. 2011). Briefly, cultures were crosslinked with 1% formaldehyde or ciprofloxacin (4 ug/mL, Topo IV only) for 20 minutes and subsequently quenched with 0.5 M glycine (formaldehyde crosslinking only). Cell pellets were collected by centrifugation and washed once with cold phosphate buffered saline (PBS). Cell pellets were resuspended with 1.5 mL of Solution A (10 mM Tris–HCl pH 8.0, 20% w/v sucrose, 50 mM NaCl, 10 mM EDTA, 10 mg/ml lysozyme, 1 mM AEBSF) and incubated at 37° C for 30 min. After incubation, 1.5 mL of 2x IP buffer (100 mM Tris pH 7.0, 10 mM EDTA, 20% triton x-100, 300 mM NaCl and 1mM AEBSF) was added and lysates were incubated on ice for 30 minutes. Lysates were then sonicated 4 times at 30% amplitude for 10 seconds of sonication and 10 seconds of rest. Lysates were pelleted by centrifugation at 8000 RPMs for 15 minutes at 4° C. Each IP was done with 1 mL of cell lysate and 40 µL was taken out prior to addition of the antibody as an input control. IPs were performed using rabbit polyclonal antibodies against DnaC (Smits et al., 2010), RNAP (Santa Cruz Biotech), GFP (Abcam, gyrase) and Myc (Invitrogen, Topo IV). IPs were rotated overnight at 4° C. After incubation with the antibody, 30 µL of 50% Protein A sepharose beads (GE) were added and IPs were incubated at RT for one hour with gentle rotation. Beads were then pelleted by centrifugation at 2000 RPM for 1 minute. The supernatant was removed and the beads were washed 6x with 1mL of 1x IP buffer. An addition wash was done with 1 mL of TE pH 8.0. After the washes, 100 µL of elution buffer I (50 mM Tris pH 8.0, 10 mM EDTA, 1% SDS) was added and beads were incubated at 65° C for 10 minutes. Beads were pelleted by centrifugation at 5000 RPMs for 1 minute. The supernatant was removed, saved and 150 µL of elution buffer II (10 mM Tris pH 8.0, 1 mM EDTA, 0.67% SDS) was added. Beads were then pelleted by centrifugation at 7000 RPMs for 1 minute and the supernatant was combined with the first elution. The combined eluates were then de-crosslinked by incubation at 65° C for overnight. The eluates were then treated with proteinase K (0.4 mg/mL) at 37° C for 2 hours. DNA was then extracted with a GeneJet PCR purification Kit (Thermo) according to the manufacturer’s instructions.

### DNA:RNA hybrid immunoprecipitation assays (DRIPs)

DRIPs were performed as described (García-Rubio et al. 2018; Lang et al. 2017; Sanz and Chédin 2019). Precultures were diluted to OD600 of 0.05 in LB and grown at 30° C with shaking. At OD600 ∼0.1, cultures were induced with 1 mM IPTG (final concentration) and grown until the culture was at OD600 = 0.3. Cells were pelleted by centrifugation and washed twice with cold PBS. Total nucleic acids were purified from cell pellets using phenol:chloroform extraction and ethanol precipitation. After drying, DNA was resuspended in TE pH 8.0 and treated with HindIII, EcoRV, EcoRI, DraII, and PstI overnight at 37° C. Digested chromosomal DNA was then purified by phenol:chloroform extraction and brought to final volume of 125 µL. Nucleic acids were then quantified using a Qubit (Invitrogen) and 10 μg were added to each IP in 470 total µL of TE. 20 µL was then removed kept as INPUT. 51 µL of 10x Binding buffer (100 mM NaPO_4_ pH 7.0, 1.4 M NaCl, 0.5% Triton X-100) was added. S9.6 antibody (Millipore) was added and samples were incubated overnight at 4° C with gentle rotation. After incubation with the antibody, 40 µL of 50% Protein A sepharose beads (GE) were added and IPs were incubated at 4° C for 2 hours with gentle rotation. Beads were then pelleted by centrifugation at 2000 RPM for 1 minute. The supernatant was removed and the beads were washed 3x with 1mL of 1x Binding buffer. After the washes, 120 µL of elution buffer of elution buffer II (10 mM Tris pH 8.0, 1 mM EDTA, 0.67% SDS) and 7 µL Proteinase K (Qiagen) was added. For the INPUT samples, 27 µL of TE pH 8 and 3 µL Proteinase K was added. All samples were incubated at 55° C for 45 minutes. Beads were then pelleted by centrifugation at 7000 RPMs for 1 minute and the supernatant moved to a new tube. DNA was purified by phenol:chloroform extraction and ethanol precipitation and used to prepare Illumina libraries using the Nextera NT library prep kit (Illumina) or analyzed using qPCR. DRIP-qPCR analysis was done by the ratio of signal at the conflict region (primer pair 188/189) divided by a control locus *yhaX* (192/193).

### RNA extraction and cDNA preparation

For assays with transcriptional reporters, cells were grown in LB to mid-exponential phase and backdiluted to OD600 0.05 into LB either supplemented with or lacking 1mM IPTG. For exponential phase experiments, cells were grown for 2 hours at 30° C (3 generations) prior to harvesting. 5 mL of culture was harvested by addition to an equal volume of ice-cold methanol followed by centrifugation at 4,000xg for 5 minutes. Cells were lysed with 20 µg/mL lysozyme for 10 minutes for cultures grown to exponential phase. RNA was isolated with the GeneJET RNA Purification Kit (ThermoFisher Scientific). 1 µg of RNA was treated with RNase-free DNase I (ThermoFisher Scientific) for 40 minutes at 37°C. DNase I was denatured by the addition of 1ul of EDTA and incubation at 65°C for 10 minutes. Reverse transcription was performed with iScript Supermix (BioRad) as per manufacturer’s instructions. mRNA abundance was measured via qPCR analysis by measuring the signal ratio of the target locus *lacZ* (primer pair 188/189) by the control *rrn* locus (primer pair 86/87).

### Next generation sequence analysis

Sequencing libraries were generated using the Nextera NT library prep kit from Illumina. Approximately 4M x 150 bp paired-end Illumina Next-Seq reads per sample were mapped to the genome of *B. subtilis* strains HM1300 (head-on lacZ) and HM1416 (co-directional lacZ) in the strain background JH642 (GenBank: CP007800.1) using Bowtie 2 (Langmead and Salzberg, 2012). Both PCR and optical duplicates were removed using Picard v1.3. Bam files were normalized for the total number of reads and the ratio of the immunoprecipitation versus the input was done using deepTools (Ramirez et al. 2014). Plots were generated in IGV (Robinson et al. 2011). Duplicate experiments were conducted for each ChIP and DRIP sequencing experiment; representative plots are shown in the figures.

## Data availability

Deep sequencing data will be uploaded to publically accessible NCBI databases prior to publication.

